# evedesign: accessible biosequence design with a unified framework

**DOI:** 10.64898/2026.03.17.712115

**Authors:** Thomas A. Hopf, Artem Gazizov, Sergio Garcia Busto, Ethan Eschbach, SunJae Lee, Milot Mirdita, Rose Orenbuch, Khaoula Belahsen, David Ross, Chris Sander, Martin Steinegger, Simon d’Oelsnitz, Debora S. Marks

## Abstract

Machine learning methods for protein engineering are rarely interoperable, require bespoke workflows, and remain inaccessible to non-experts. Yet the design problems that matter most – conditional design subject to real-world constraints, multi-objective optimization, and iterative lab-in-the-loop workflows where experimental data continuously refines successive design rounds – demand exactly the kind of flexible, composable infrastructure that no single tool provides. We present evedesign, a unified open-source framework that formalizes conditional biosequence design in a method-agnostic way, enabling complex multiobjective workflows combining supervised and unsupervised models from standardized specifications, and built from the outset to support iterative experimental integration. An interactive web interface facilitates end-to-end design for a broad scientific audience at https://evedesign.bio. We demonstrate evedesign’s utility in antibody engineering, enzyme design, and natural enzyme discovery, and invite open-source community contributions.

## Introduction

Protein engineering has the potential to address critical unmet needs in next-generation therapeutics, sustainable biomanufacturing, biosecurity, pollutant remediation, and climate adaptation^1^. Recent years have seen an extraordinary proliferation of machine learning methods for protein design, spanning MSA-based approaches that train on evolutionary sequences^2–11^, large language models trained on unaligned protein sequences^12–15^, inverse folding models^16,17^, and *de novo* 3D structure design models^18–21^. Despite impressive experimental demonstrations across this landscape, translating these methods into real-world design workflows remains difficult even for experts and continue to require bespoke, labor-intensive code development.

Two interconnected gaps limit the practical impact of this progress (Table 1). First, the absence of a standardized programming interface means that methods cannot be readily compared, combined, or swapped. This lack of interoperability is especially costly for conditional design – where proteins are engineered subject to real-world constraints such as thermostability, pH tolerance, removal of T cell epitopes, or binding to a new target – and for iterative lab-in-the-loop workflows that incorporate new experimental data to guide successive design rounds. These multi-objective, constraint-driven tasks are at the heart of most real-world design projects, yet existing frameworks are either not model-agnostic or fail to generalize beyond sequence-based models for monomeric proteins^21–26^. The lack of interoperability extends to the controlled integration of models into agentic systems^27–31^. Second, there is no open-source, interactive user interface that spans the complete design workflow from target protein to orderable DNA sequences, while remaining applicable independent of the underlying models and pipelines^32–35^. Proprietary solutions exist^36–41^, but the lack of an open alternative forces non-computational researchers to rely on disconnected tools and limits reproducibility and accessibility across the community.

**Table 1.** Comparison of evedesign with single-method tools and ad hoc workflows across key design capabilities.

| Capability | Single-method tools / ad hoc workflows | evedesign |
| --- | --- | --- |
| Model interoperability | Each tool has its own API and data format | Standardized interface: any model can be swapped or combined |
| Design representation | Fixed single-level (sequence or structure or embedding) | Multi-level: sequence, embedding, and 3D structure simultaneously |
| Conditional design | Hardcoded per tool | Declarative system specification; constraints composable at runtime |
| Multi-objective optimization | Requires bespoke glue code for each combination | Native: compose score/generate/transform operations across models |
| Supervised + unsupervised integration | Manual, ad hoc | First-class support: attach experimental labels directly to instances |
| Lab-in-the-loop iteration | Not supported | Designed for it: declarative, serializable pipelines; halt, replay, and update with new data |
| User interface | None, or tool-specific | End-to-end: target sequence to orderable DNA, model-agnostic |
| Privacy & self-hosting | Varies; often unavailable | *evedesign_server* can be hosted on private infrastructure |
| Access & reproducibility | Proprietary or fragmented | Open-source, FAIR-compliant, self-hostable |

To address these gaps and bring protein design into alignment with FAIR principles^42^ (findability, accessibility, interoperability, and reusability), we introduce evedesign: a multi-layered open framework comprising (i) a standardized model interface with reference implementations in a Python package, (ii) a REST API and pipeline runner, for executing design tasks from declarative job specifications, and (iii) an interactive user interface covering complete design workflows, publicly available at https://evedesign.bio. The framework is designed as a living community resource: its modular architecture enables new models and workflows to be incorporated as the field advances, ensuring that evedesign remains a current and extensible solution for both computational and experimental researchers.

## Results and Discussion

### A unified interface for biomolecular design

#### Framing design as a conditional modeling problem

A central obstacle to applying diverse protein design methods in practice is that each operates on its own representation of the biological problem (sequences, structures, embeddings, or alignments) with no common language between them. To resolve this, evedesign frames biomolecular design as a conditional modeling problem (Figure 1A): the user specifies a molecular system composed of individual entities (protein, DNA or RNA chains, or ligands), and any known information about each entity, such as primary sequence, 3D structure, homologs, binding partners, post-translational modifications, or multimeric state, is supplied as a standardized declarative data structure. Sequences can be fully specified, partially masked, or left open for design, and structures can include fragments or alternative conformational states. This mirrors the conditioning capabilities now emerging in next-generation structure prediction and design methods^20,43^, but generalizes them to arbitrary combinations of models and design objectives (full specification in Supplementary Figure 1).

**Figure 1:**
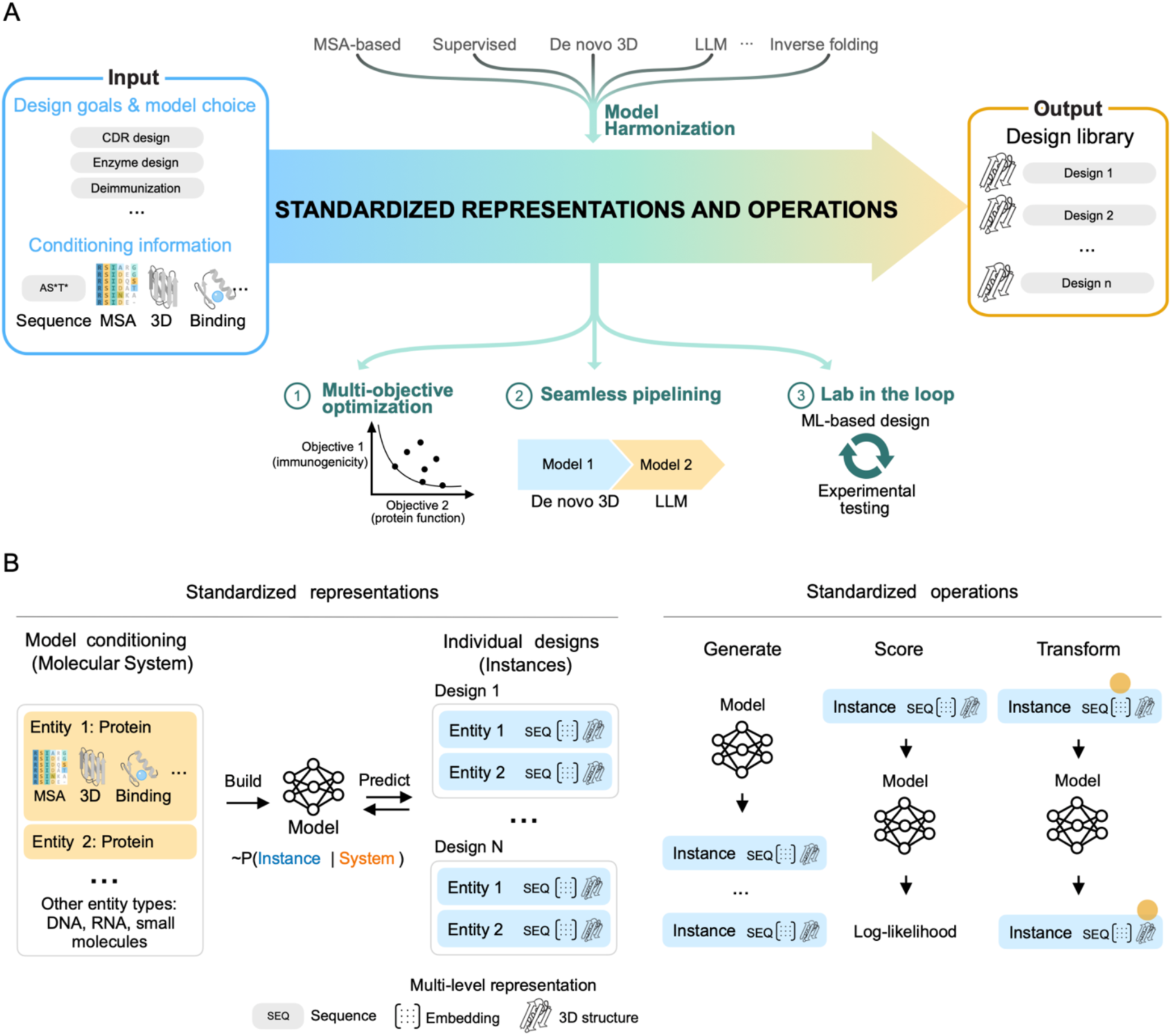
Unified framework for biomolecular design. **(A)** evedesign enables key biomolecular design applications by unifying different method types that are otherwise not interoperable without bespoke development. Design pipelines and their user inputs are specified in a method-agnostic way and executed through standardized operations, facilitating the creation of complex multi-objective optimization and lab-in-the-loop design workflows. **(B)** *Left*: We frame biomolecular design as a conditional modeling problem where models are first set up on a molecular system composed of protein, nucleotide or small molecule entities with any known information. Conditioned models generate or predict individual designs ("instances") which carry a per-entity multi-level representation that allows designs to flow seamlessly between methods operating on different input types. *Right*: Design workflows are expressed by combining three standardized model operations. Because all operations share the standardized instance format, outputs from one model are directly consumable as inputs to another without reformatting.

The overall design goal, for example, optimizing thermostability while removing T cell epitopes, emerges naturally from the user’s choice of models applied and the system-level constraints supplied to the models, rather than being explicitly declared.

#### A multi-level instance representation

Once set up for a particular conditional modeling problem, a model operates on instances: concrete realizations of the molecular system, such as candidate mutants or newly generated sequences. Each instance carries a multi-level representation encoding information at the sequence, positional embedding, and 3D structure (PDB format) levels simultaneously, though not all levels need to be populated (Figure 1B). This layered scheme, in contrast to frameworks that enforce a single fixed representation, enables designs to flow seamlessly between methods operating on different input types. For example, a single pipeline can score designs using both sequence likelihoods and 3D structural constraints, map LLM-derived embeddings to coordinates for structural validation, or leave coordinates unspecified for flexible regions handled by sequence-only models.

The instance format also supports supervised learning workflows for lab-in-the-loop applications: experimental measurements can be attached as labels to instances, and embeddings or likelihoods from unsupervised models used as input features to train regression models predicting user-defined properties of interest.

#### Three composable operations: generate, score, transform

To cover the full range of design and prediction tasks without bespoke code, we propose that all workflows can be expressed through three standardized model operations (Figure 1B), semantically analogous to general machine learning frameworks such as scikit-learn^44^ but adapted for the multi-level instance representation:

- *Generate* produces new design instances from the model, conditioned on the system specification, at the model’s preferred representation level(s).
- *Score* assigns a quantitative fitness value, preferably a log-likelihood, to each instance, enabling comparison across designs or mutants.
- *Transform* maps instances between representation levels; for example, predicting 3D structure from sequence, or computing embeddings from sequence.

Because all operations share the standardized instance format, outputs from one model are directly consumable as inputs to another without reformatting. This allows complex, multi-objective workflows to be assembled by composition: for instance, Gibbs sampling over a sequence space guided by multiple scoring models, each of which internally transforms sequences to embeddings via an LLM before applying a supervised predictor.

#### The evedesign software packages

Our framework is built modularly to serve a broad set of use cases and user communities. The core MIT-licensed Python package *evedesign* implements the interface above and currently includes reference implementations of three key model classes: (i) a new evolutionary method, EVmutation2 (described below); (ii) the ESM-2 protein language model^14^; and (iii) ProteinMPNN/LigandMPNN for inverse folding^16,45^. These are bundled with a general-purpose multi-entity Gibbs sampler, sequence distance restraints^10^, utilities for MSA generation with MMseqs2^46^, 3D structure template search and mapping with FoldSeek^47,48^, sequence space analysis, and codon optimization^49^. We invite the community to contribute implementations of additional models under the shared specification.

On top of the core package, the MIT-licensed Python package *evedesign_server* implements a REST API and lightweight design pipeline execution engine package for distributed execution and status tracking of design jobs (Supplementary Figure 2). This package allows hosting a full design server backend on the user’s own computing infrastructure for commercial and privacy-sensitive deployments.

Our React-based frontend application provides a user-friendly way to run design jobs end-to-end from target sequence to codon-optimized nucleotide sequences without writing code or executing tools on the command line. After job completion, the user interface allows interactive exploration of the generated designs in the context of available 3D structures and evolutionary sequences, with rich data visualizations (Figure 2).

**Figure 2:**
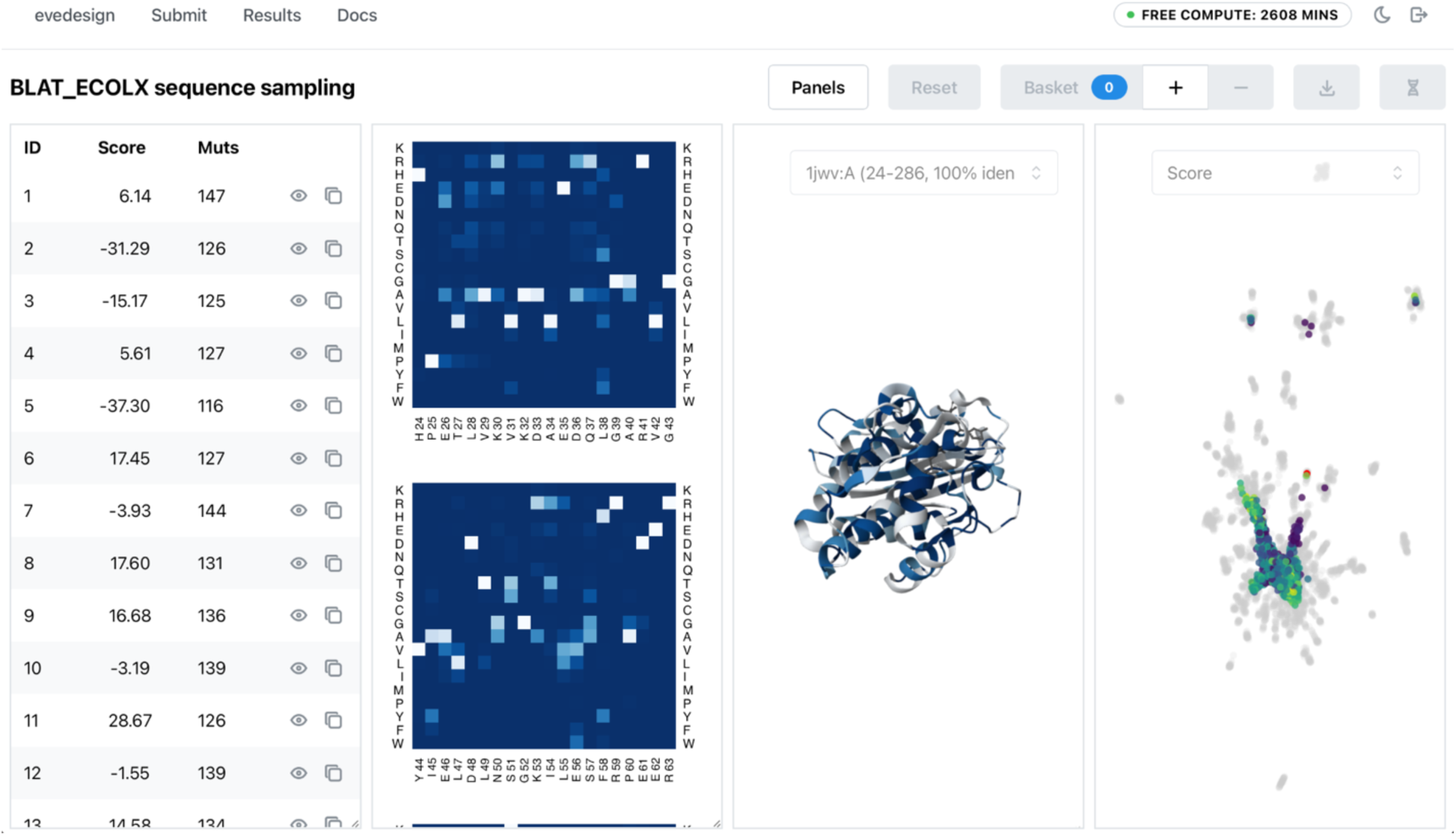
Interactive user interface for biomolecular design. A web-based user interface allows to submit design jobs and analyze and filter the generated sequences interactively upon job completion through multiple linked visualizations, including a list of designs (1st panel from left), marginal amino acid frequencies (2nd panel), in the context of 3D structures (3rd panel) and the sequence space of natural sequences (4th panel). After curation, the final sequence selection can be forwarded to DNA sequence codon optimization and exported for ordering at a nucleotide synthesis provider.

#### EVmutation2: a lightweight evolutionary model for generative design

Evolutionary models have a strong experimental track record for designing functional proteins far from their natural sequence^9,10,50^, but existing approaches including our own EVE^4^ require training or fine-tuning on individual MSAs, making them too slow for use in an interactive server. We therefore developed EVmutation2, a new MSA-based evolutionary model designed specifically for fast generative inference. EVmutation2 couples an order-invariant autoregressive decoder to the single- and pair-sequence representations of a compact AlphaFold3^43^ reimplementation (14.3M parameters), going beyond BERT-style masked prediction to enable direct sequence sampling^6^. All known tokens are moved to the prefix in randomized order, maximizing context for designed positions regardless of their location in the sequence^16^. Decoder attention weights are derived from the pair representation by linear projection rather than full query-key-value attention^51^, enforcing biologically meaningful pairwise interactions while reducing parameter count (see Supplementary Methods). The model was trained on the OpenProteinSet^52^ to reconstruct randomly masked MSA sequences via cross-entropy loss. Despite its compact size, EVmutation2 performs on par with EVE on ProteinGym benchmarks (Supplementary Figure 3) while requiring no per-target fine-tuning, making it well suited to a general-purpose community server.

### Case studies: demonstrating evedesign workflows

The following case studies are not intended to benchmark model performance or claim state-of-the-art results. Rather, they illustrate how the standardized interface, composable operations, and multi-level instance representation of evedesign enable diverse real-world design tasks – unsupervised generative design, structure- and sequence-based optimization, and supervised property prediction – to be expressed as coherent, reproducible workflows without bespoke code.

#### Unsupervised enzyme design with an evolutionary model

To demonstrate generative sequence design using the evolutionary model introduced above, we reproduced a workflow analogous to Russ *et al.*^9^, who used a Potts model to design diversified functional variants of chorismate mutase (EcCM), a key enzyme in aromatic amino acid biosynthesis, validated by high-throughput in vivo complementation assay.

Starting from an MMseqs2-derived MSA of natural EcCM homologs, we built an EVmutation2 model and confirmed that its zero-shot scores discriminate functional from non-functional sequences in both the natural homolog set and the Russ *et al.* experimental designs (AUROC = 0.77 and 0.81 respectively; Figure 3A; Supplementary Figure 4), establishing that the model captures functional constraints in this family.

**Figure 3:**
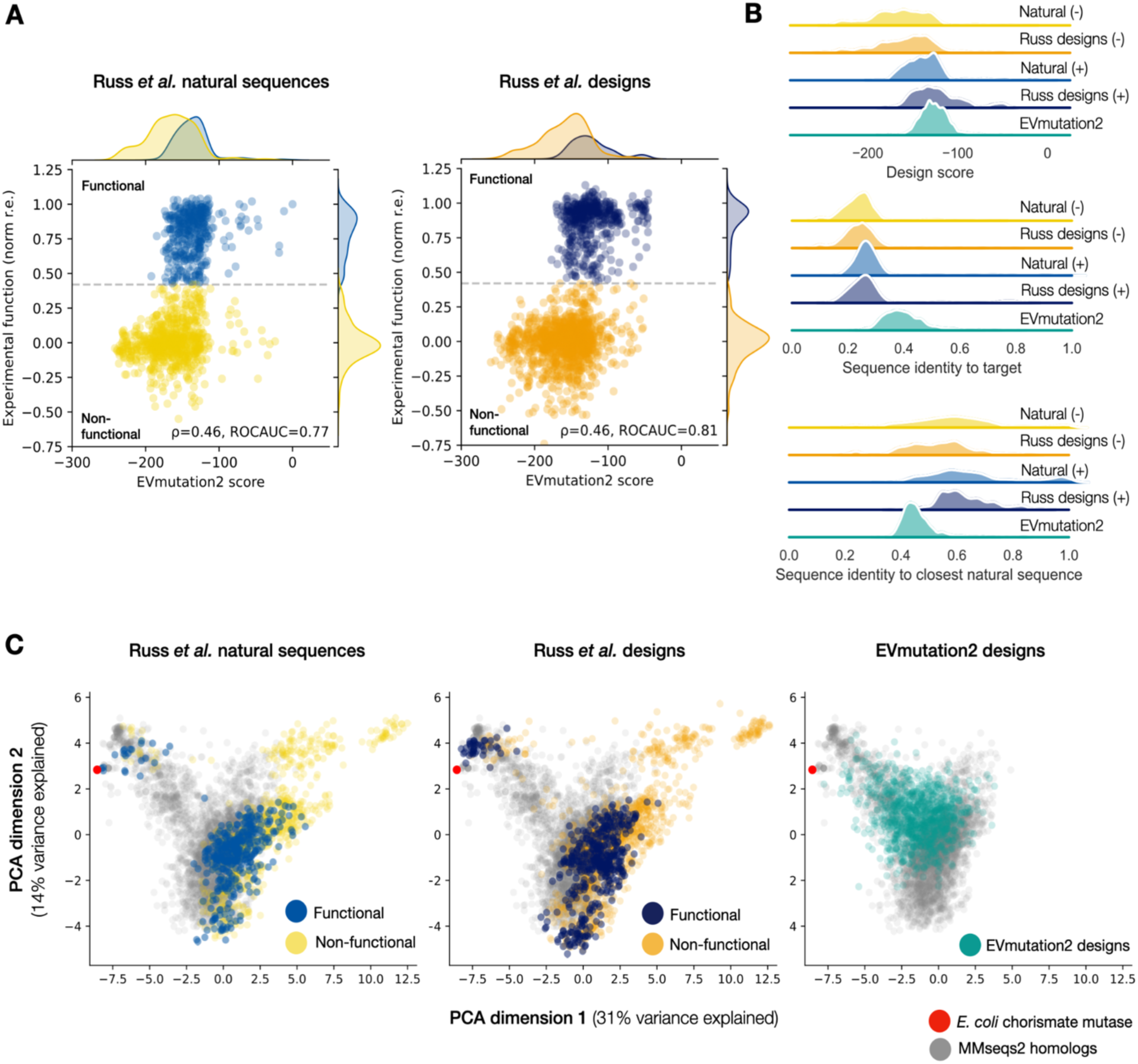
Unsupervised enzyme design with an evolutionary sequence model. **(A)** Zero-shot scores from EVmutation2 (x-axis) discriminate between functional and non-functional chorismate mutase sequences (y-axis, normalized relative enrichment > 0.42 for functional sequences as in Russ *et al.*). *Left*: natural homologs, *right:* computationally designed sequences. **(B)** *Top*: Designs generated from EVmutation2 have comparable zero-shot scores to functional chorismate mutase sequences (+) and are assigned more favorable scores than non-functional sequences (-). *Middle*: EVmutation2 designs are more similar to *E. coli* chorismate mutase than sequences from Russ *et al. Bottom*: EVmutation2 designs are more distant to the closest natural sequence than Russ *et al.* design. **(C)** Projection of natural and designed sequences with principal component analysis indicates that Russ *et al.* sequences inhabit a distinct, partially overlapping region of sequence space compared to our natural sequences identified with MMseqs2 and designs from EVmutation2.

We then used evedesign’s *generate* operation to produce 1,024 new sequences by autoregressive sampling. The designs recapitulate the first- and second-order amino acid statistics of natural sequences (*r* = 0.98 and *r* = 0.96; Supplementary Figure 5), indicating that coevolutionary constraints are preserved. Their log-likelihood scores are comparable to functional natural sequences and Russ *et al.* designs, and substantially higher than non-functional sequences (Figure 3B). In sequence space, EVmutation2 designs interpolate the MMseqs2-defined natural sequence distribution and are on average more similar to EcCM than Russ *et al.* designs (median identity 39% vs. 25%), while still showing meaningful extrapolation from the closest natural sequence (Figure 3C). Experimental validation of these designs is beyond the scope of this study, but the workflow demonstrates how a fully unsupervised generative pipeline can be assembled and executed within evedesign in a few lines of code.

#### Complementary sequence- and structure-based antibody scoring

To illustrate how evedesign enables sequence and structure models to be combined within a single workflow, we scored single-residue substitutions across the variable chains of seven clinically relevant antibodies previously studied by Hie *et al.*^53^, who used ESM-1 ensembles for language model-guided affinity maturation.

Applying evedesign’s *score* operation with ESM-2, we found that 44 of 54 experimentally tested beneficial mutations (81.5%) ranked in the top 5% of ESM-2 scores (Figure 4A, Supplementary Figure 6), consistent with the original findings. Notably, ESM-2 assigned lower scores to light chain mutations that decreased or did not affect antigen binding (*p* = 0.03, Mann-Whitney U-test), though this effect was not significant for heavy chain mutations, a distinction not captured by the original ESM-1 ensemble.

**Figure 4:**
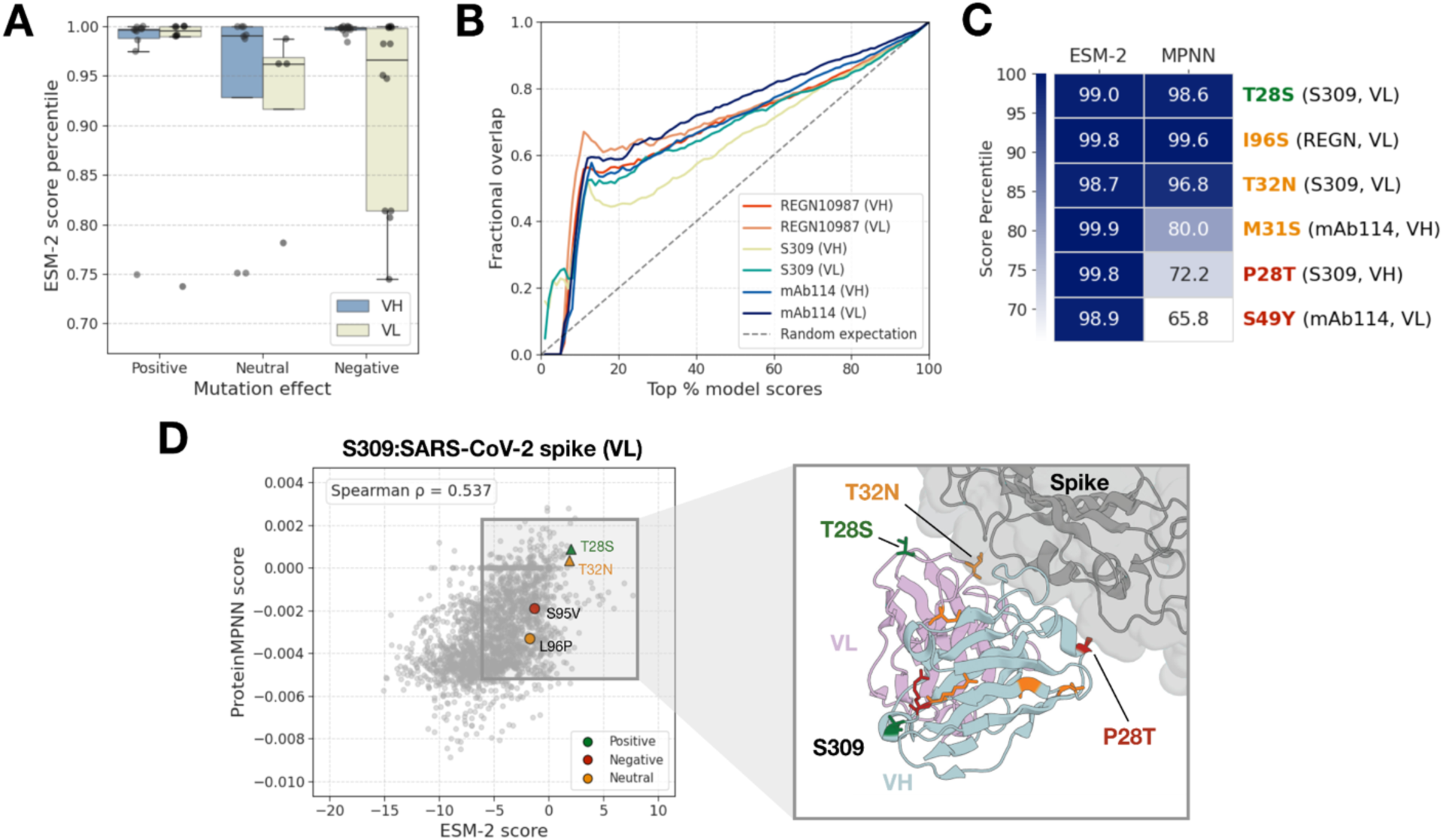
Structure- and sequence-based zero-shot antibody maturation. **(A)** ESM-2 score percentiles of single substitution mutation effects that were experimentally characterized in Hie *et al.*, for different variable chains (heavy, VH; light, VL). For VL, neutral and negative mutants have significantly lower scores than positive mutants (*p* = 0.03, Mann-Whitney U-test). **(B)** Fractional overlap between the top ProteinMPNN and ESM-2 model scores as a function of top model scores for the three antibody-antigen complexes with available structures (PDB IDs: 6WPS, 6XDG, 5FHC). The low overlap within highly-ranked mutants highlights the opportunity to combine structure- and sequence-based models for the discovery of antibody variants with increased affinity. **(C)** ProteinMPNN and ESM-2 score percentiles for affinity-decreasing (red), affinity-increasing (green) and neutral (orange) single mutations at sites near antibody-antigen interfaces for the three complexes considered. While ESM-2 predicts all mutant classes to be favorable, ProteinMPNN can downscore neutral and deleterious substitutions. **(D)** Key trends at antibody-antigen interfaces seen in experimental data from Hie *et al.* are captured by ProteinMPNN. *Left*: ProteinMPNN and ESM-2 scores for all single substitutions for the light chain of the S309 antibody in complex with the SARS-CoV-2 spike protein. Experimentally characterized mutations by Hie *et al.* are shown colored according to the mutation effect, and mutants near the antibody-antigen interface are highlighted with a triangle. *Right*: the S309:SARS-CoV-2 complex structure (PDB ID: 6WPS) with mutations at sites near the antibody-antigen interface shown.

Crucially, because unsupervised language models are trained on sequences without structural context, they are not expected to explicitly capture binding interface effects. For the three antibody-antigen complexes with available experimentally determined structures^54–56^, we additionally applied evedesign’s *score* operation with ProteinMPNN, which conditions on 3D coordinates. ProteinMPNN and ESM-2 scores were globally correlated but showed minimal overlap in their highest-scoring mutations (mean top-5% overlap = 0.08 ± 0.11; Figure 4B, Supplementary Table 1). Importantly, ProteinMPNN assigned substantially lower scores to deleterious and neutral mutations at interface-proximal positions (within 6Å of the antigen), while correctly ranking highly the only beneficial interface mutation across all complexes (Figure 4C, D). This divergence illustrates a key advantage of the multi-model evedesign framework: sequence- and structure-based scores provide complementary information and can be straightforwardly combined to prioritize mutations, particularly for interaction-sensitive applications such as antibody maturation.

#### Supervised discovery of efficient enzyme variants

To demonstrate supervised property prediction within evedesign, we reproduced an analysis from Eom *et al.*^57^, who trained a Random Forest regressor on ESM-1b embeddings of a curated kynureninase (KYNase) kinetics dataset to mine natural sequence databases for homologs with improved catalytic efficiency.

Using evedesign’s *transform* operation to compute ESM-2 embeddings and its instance labelling scheme to attach experimental kcat/Km values as training targets, we trained an equivalent regressor and recovered cross-validation performance matching the original study (Spearman *ρ* = 0.73 vs. 0.72; Figure 5A, B; Supplementary Figure 7). Ranking of 5,676 unlabelled homologs with the *score* operation reproduced the key experimental outcome: the highest-efficiency variant (K3) ranked consistently in the top 10 across random seeds, while lower-efficiency variants ranked progressively lower (Figure 5C; Supplementary Tables 2–3; Supplementary Figure 8). This confirms that evedesign’s supervised workflow generalizes to database-scale property-guided search with no modification to the underlying framework.

**Figure 5:**
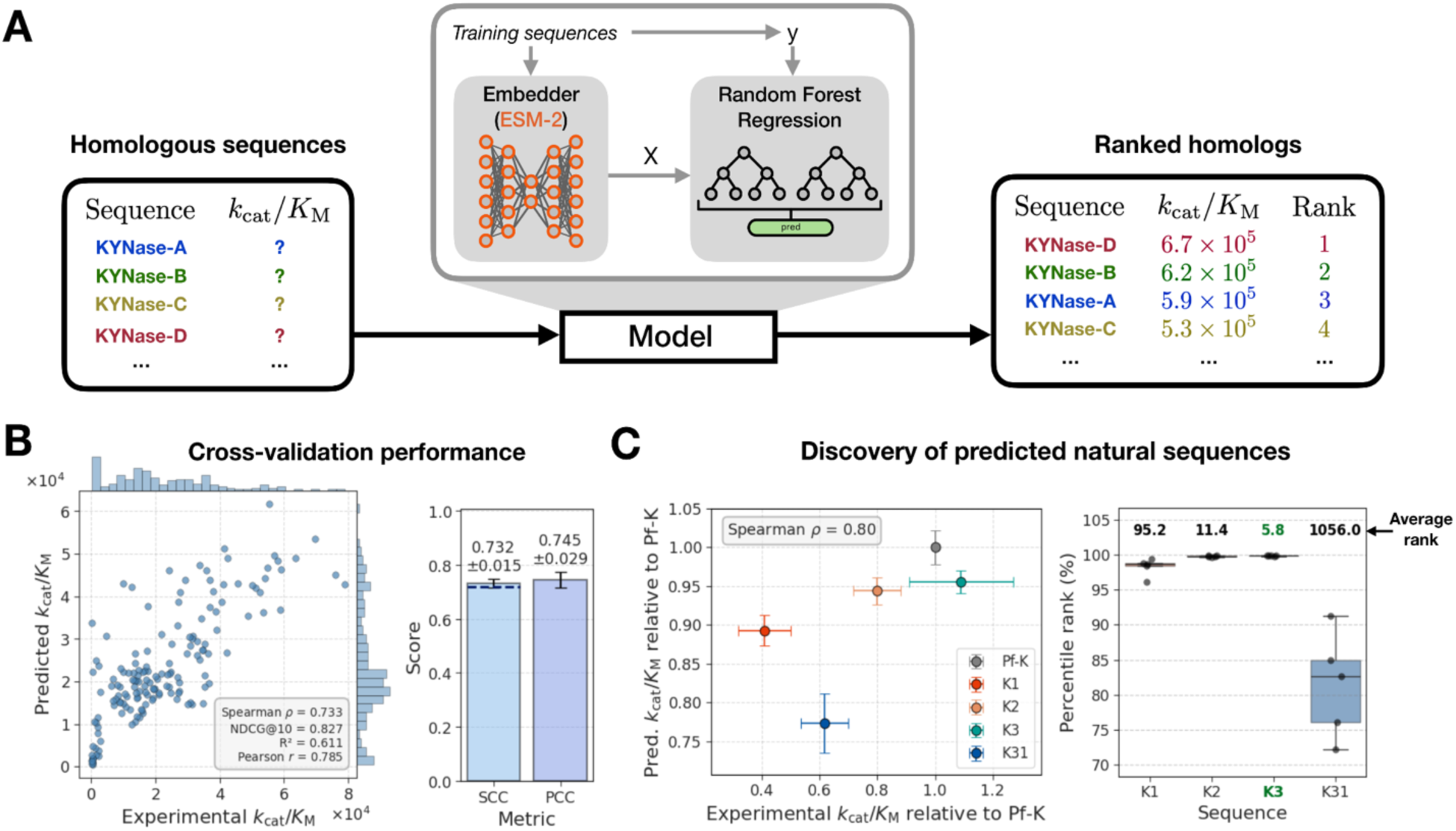
Supervised discovery of efficient KYNases. **(A)** The supervised pipeline to predict catalytic efficiencies using Random Forest regression on ESM-2 embeddings. Unlabelled homologs are scored and ranked using the trained model. **(B)** Five-fold cross-validation catalytic efficiency predictions and performance for one random initialization (*left*) and five independent initializations (*right*). Data was split randomly and scikit-learn default hyperparameters were used for the Random Forest. The blue dashed line indicates the Spearman Correlation Coefficient (SCC) reported by Eom *et al.*, showing that performance is comparable to that of the original study using ESM-2. The Pearson Correlation Coefficient (PCC) is also shown, and error bars represent standard deviations. **(C)** *Left*: experimental measurements of KYNases from Eom *et al.* and predicted catalytic efficiencies using the evedesign pipeline across five different initializations, which together highlight a broad agreement of predictions with experimental data. All values are relative to the starting Pf-K sequence (gray), and error bars represent standard deviations. *Right*: percentile of predicted ranked efficiencies by evedesign, alongside the average absolute rank out of 5,676, for the top three (K1-3) and 31st (K31) highest-ranked homologs from Eom *et al.* KYNase, kynureninase; Pf-K, *Pseudomonas fluorescens*.

## Conclusion

We have introduced evedesign, a unified open-source framework that addresses the fragmentation and accessibility barriers currently limiting the real-world impact of machine learning methods for protein engineering. By standardizing how models represent molecular systems, exchange designs, and compose into pipelines, evedesign makes it practical to tackle the conditional, multi-objective design problems that matter most in therapeutic development, biosecurity, and sustainable biomanufacturing, without requiring bespoke implementation for each new task. We demonstrated this across three case studies spanning unsupervised generative design, complementary sequence- and structure-based scoring, and supervised property-guided discovery.

The modular architecture is intentionally designed to serve a broad community: computational biologists and ML researchers can integrate new models by implementing a small set of Python interface methods and retaining full control of their own code, while experimentalists can access the same workflows through the interactive interface at https://evedesign.bio. For users with data privacy requirements, including those in commercial or clinical settings, the *evedesign_server* package can be self-hosted on private infrastructure, ensuring that proprietary sequences and experimental data never leave the user’s own computing environment. As a FAIR-compliant, MIT/AGPLv3-licensed platform, evedesign provides the transparency that rigorous ML-guided design requires.

The framework presented here is a living software foundation rather than a finished product. We plan to extend the model library, including *de novo* structure design methods, and we note that the pipeline runner’s declarative job specification and serializable instance format were designed from the outset to support fully iterative lab-in-the-loop workflows, where new experimental data continuously refines the design cycle. This is a capability that ad hoc collections of disconnected tools cannot readily accommodate, and we view its full realization as the most important near-term frontier for the field. We welcome and actively invite community contributions to grow evedesign alongside the methods it integrates.

## Code and webserver availability

Our interactive user interface is available at https://evedesign.bio and provides free design computations for non-commercial users. For users requiring data privacy, *evedesign_server* can be self-hosted on private infrastructure. All source code is made available under open-source licenses (MIT for the core package, *evedesign_server* and EVmutation2; AGPLv3 for the web interface) at https://evedesign.bio/code with documentation included in all code repositories.

## Acknowledgements

This work was supported by the National Institute of Standards and Technology (award number 70NANB24H116). The authors thank Anthony Gitter for valuable comments on the manuscript.

## Supplementary Methods

### EVmutation2 model architecture

The key innovation of this model is to couple an order-invariant autoregressive decoder to the single and pair representations of AlphaFold3 that moves all known tokens to the prefix in randomized order, providing maximum context for designed positions independently of where they are located in the sequence and allowing sampling of score distributions from the model. Decoder attention weights for self- and cross-attention are derived by linear projection of the pair representation:

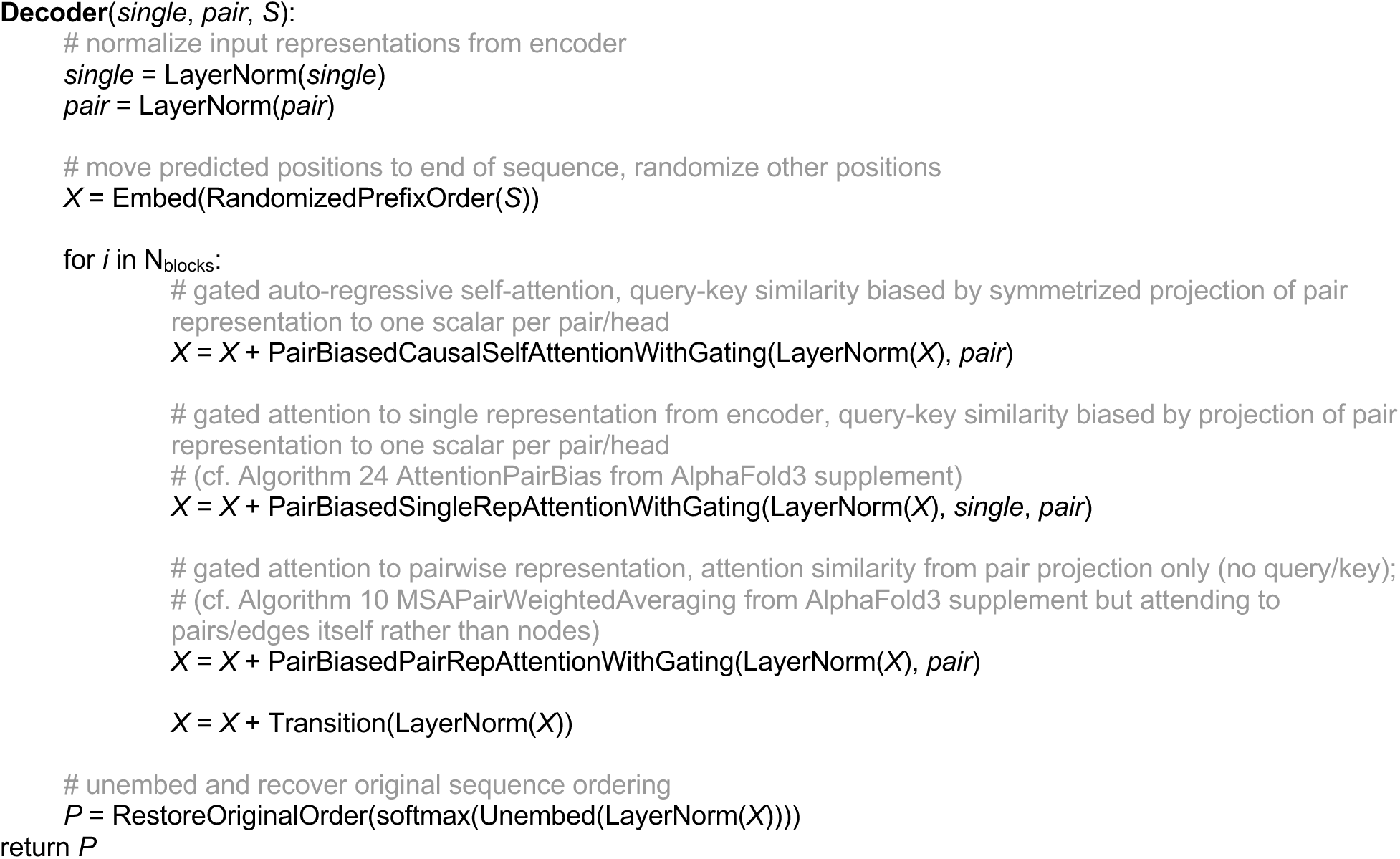

The encoder part of the model is based on an open-source reimplementation of AlphaFold3 simplified to single-chain proteins (https://codeberg.org/lucidrains/alphafold3-pytorch) and with atom-level features removed.

We trained the model from scratch for ∼10^5^ steps with the Adam optimizer and a batch size of 16. We used a base learning rate of 3 * 10^-4^, which was linearly ramped up from 0 in the first 1000 steps, and then decreased by a factor of 0.95 every 5 * 10^4^ steps as in the original AlphaFold3 schedule^43^.

The following block numbers and dimensionalities were applied to the model architecture for a small overall model size (total trainable model parameters: 14.3M):

- AlphaFold3-based encoder (10.1M parameters):

- Single representation dimensionality: 192
- Pairwise representation: dimensionality: 64
- 4 MSA module blocks
- 12 Pairformer blocks
- Autoregressive decoder (4.2M parameters)

- Single representation dimensionality: 192
- 6 decoder blocks

## Supplementary Tables

**Supplementary Table 1.** Spearman correlations between ESM-2 and ProteinMPNN model scores for three antibody-antigen complexes with experimental structures^54–56^.

| Antibody | REGN10987 |  | mAb114 |  | S309 |  |
| --- | --- | --- | --- | --- | --- | --- |
| Chain | H | L | H | L | H | L |
| Spearman $\rho$ | 0.525 | 0.529 | 0.550 | 0.673 | 0.414 | 0.537 |

**Supplementary Table 2.** Ranks and normalized scores from the supervised evedesign pipeline for the experimentally characterized KYNase homologs in Eom *et al*.^57^ A total of 5,676 homologs were evaluated. 14 homologs were discarded from the original homolog list due to being longer than the default recommended maximum sequence length of ESM-2. Scores were normalized by dividing the efficiency prediction by the maximum predicted value.

| Random seed # | K1 |  | K2 |  | K3 |  | K31 |  |
| --- | --- | --- | --- | --- | --- | --- | --- | --- |
|  | Rank | Score | Rank | Score | Rank | Score | Rank | Score |
| 251 | 218 | 0.874 | 15 | 0.948 | 3 | 0.981 | 1583 | 0.731 |
| 229 | 75 | 0.923 | 11 | 0.972 | 7 | 0.983 | 496 | 0.854 |
| 143 | 86 | 0.933 | 16 | 0.972 | 5 | 0.987 | 857 | 0.825 |
| 755 | 66 | 0.913 | 8 | 0.969 | 4 | 0.972 | 1360 | 0.767 |
| 105 | 31 | 0.929 | 7 | 0.972 | 10 | 0.968 | 984 | 0.786 |

**Supplementary Table 3.** Average ranks and normalized scores from the supervised evedesign pipeline across five different initializations (Supplementary Table 2) of the 10 highest-ranked sequences (K1-10) from Eom *et al*.^57^ Scores were normalized by dividing the efficiency prediction by the maximum predicted value.

|  | K1 | K2 | K3 | K4 | K5 | K6 | K7 | K8 | K9 | K10 | K31 |
| --- | --- | --- | --- | --- | --- | --- | --- | --- | --- | --- | --- |
| Rank | 95.2 | 11.4 | 5.8 | 1297.4 | 1576.4 | 25.2 | 43.4 | 464.6 | 322.4 | 10.4 | 1056 |
| Score | 0.915 | 0.967 | 0.978 | 0.769 | 0.745 | 0.952 | 0.935 | 0.852 | 0.872 | 0.969 | 0.793 |

## Supplementary Figures

**Supplementary Figure 1:**
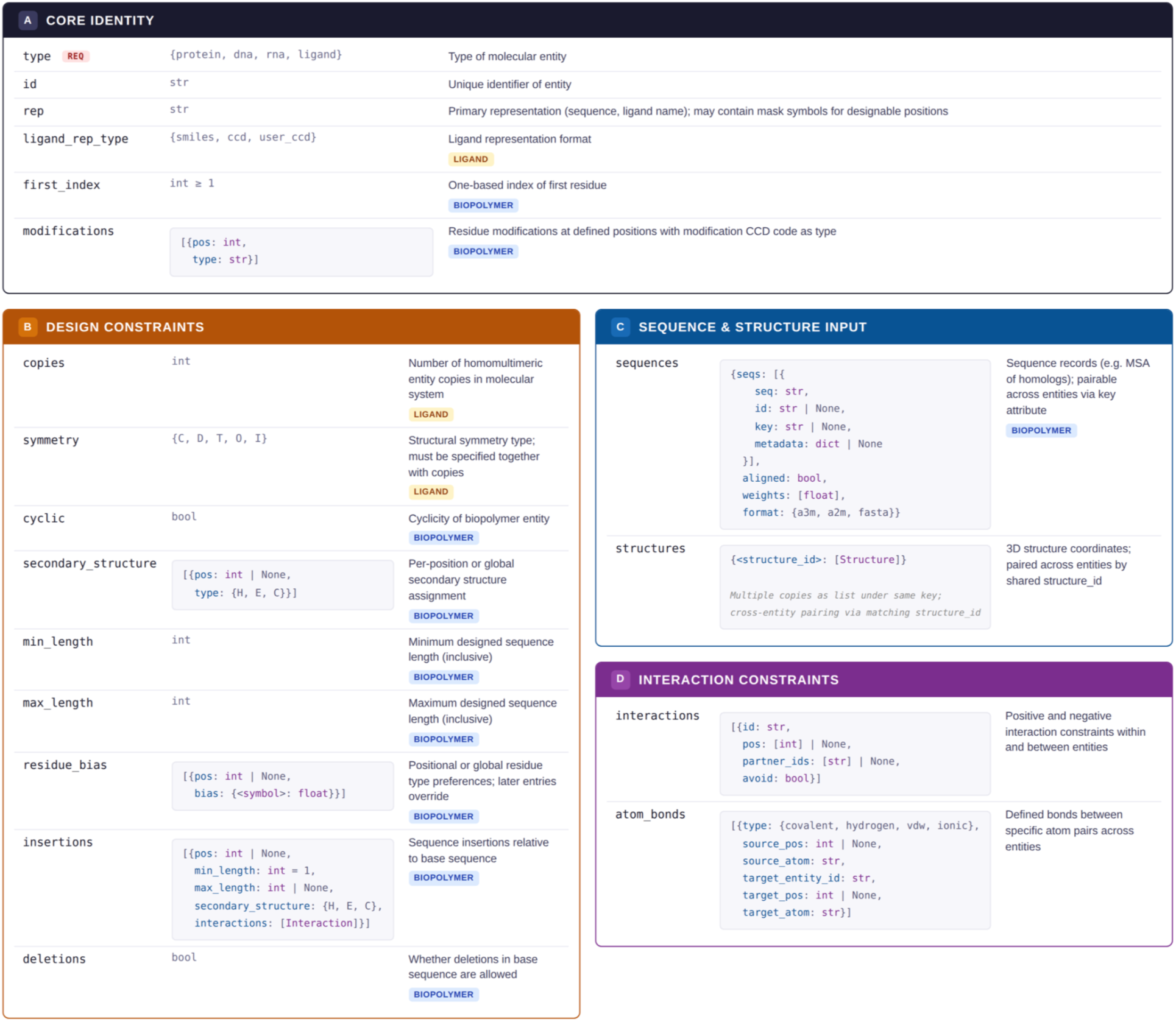
Entity attribute specification. Describing the inputs for modeling problems as a biomolecular system consisting of multiple entities allows to standardize input formats across diverse types of sequence- and structure-based models. For each protein, nucleotide or ligand entity, any known information that the modeling should be conditioned on may be supplied through various types of attributes.

**Supplementary Figure 2:**
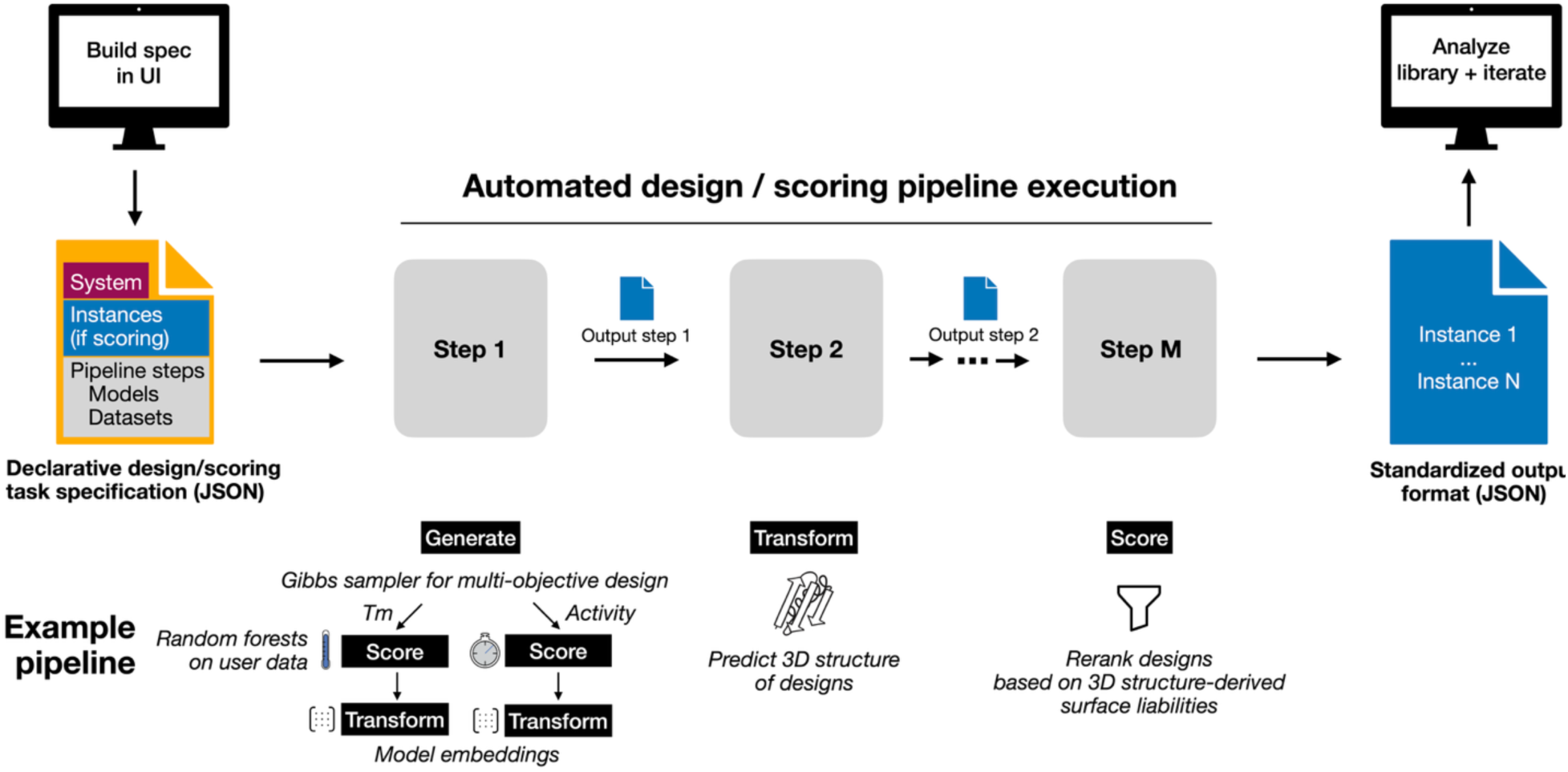
Declarative design pipelines and REST API. Expressing biomolecular design and scoring problems through our framework enables the automated execution of highly flexible, complex design pipelines from a declarative schema. At each step, a *generate*, *transform* or *score* operation is applied to create new or update the existing instances, using a single model or a composition of multiple models. While executing the pipeline, results are forwarded from step to step as a list of instances with their multi-level representations and additional metadata like scores or confidence values (blue document symbol). Our web-based interface (UI) offers a user-friendly way to specify design problems, submit the task for execution, and interactively explore the generated designs upon completion.

**Supplementary Figure 3:**
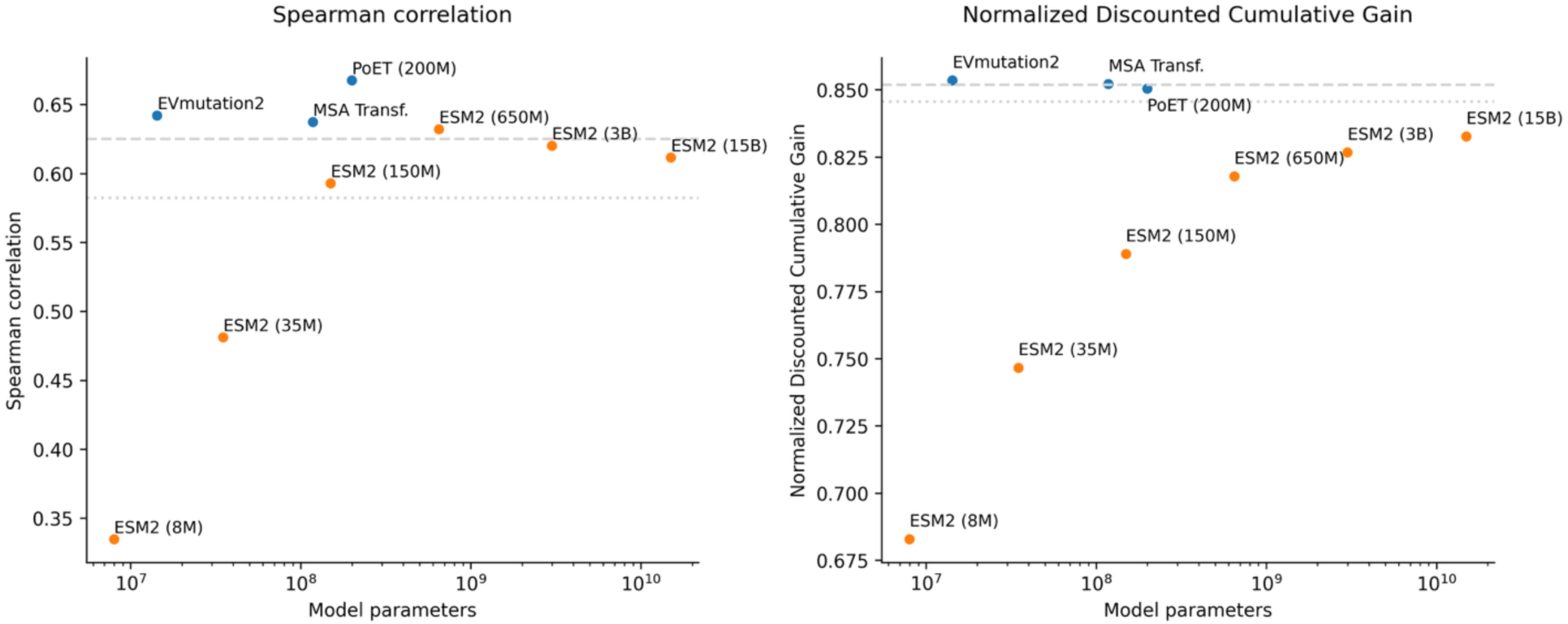
EVmutation2 evaluation on ProteinGym datasets. The performance of EVmutation2 was assessed on a curated high-quality subset of the ProteinGym^50^ zero-shot benchmark dataset by Spearman rank correlation *(left panel)* and Normalized Discounted Cumulative Gain (NDCG, *right panel*). We included a total of 32 ProteinGym datasets that achieved a Spearman correlation > 0.6 for at least one of the 74 methods available in ProteinGym at the time of curation, kept only up to 5 datasets from the same source and one dataset per target protein, and removed datasets with overly biased experimental data distributions (Supplementary Data 1). Notably, MSA-based methods (blue circles, dashed line for EVE^4^, which trains a full model per MSA) give more accurate predictions than the included baseline protein language model ESM2 at most model sizes (orange circles). EVmutation2 achieves leading results amongst the compared methods in identifying mutations at the favorable end of the distribution as measured by NDCG. We also include the performance of the first version of EVmutation^2^ for comparison (dotted line).

**Supplementary Figure 4:**
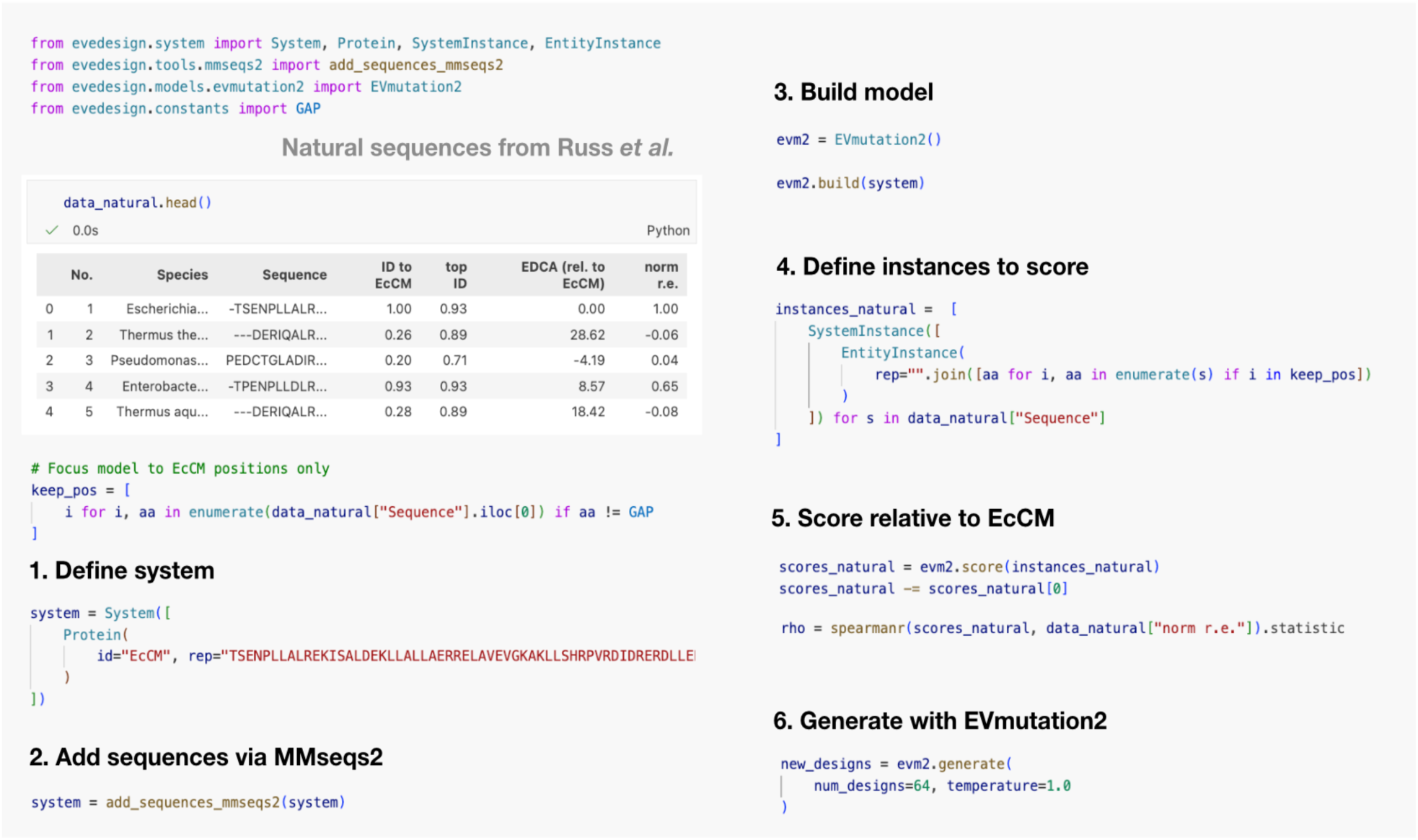
Code demonstration of the generative capabilities of evedesign. The workflow shown exemplifies the enzyme generation case study using EVmutation2, and also shows how natural sequences from Russ *et al.*^9^ can be scored. The pipeline can be summarised in six steps: defining the system, adding homologous sequences through MMseqs2, building the EVmutation2 model, defining instances to score, scoring instances and generating designs. Analogous steps can be followed for the designed sequences in Russ *et al.* to reproduce all results.

**Supplementary Figure 5:**
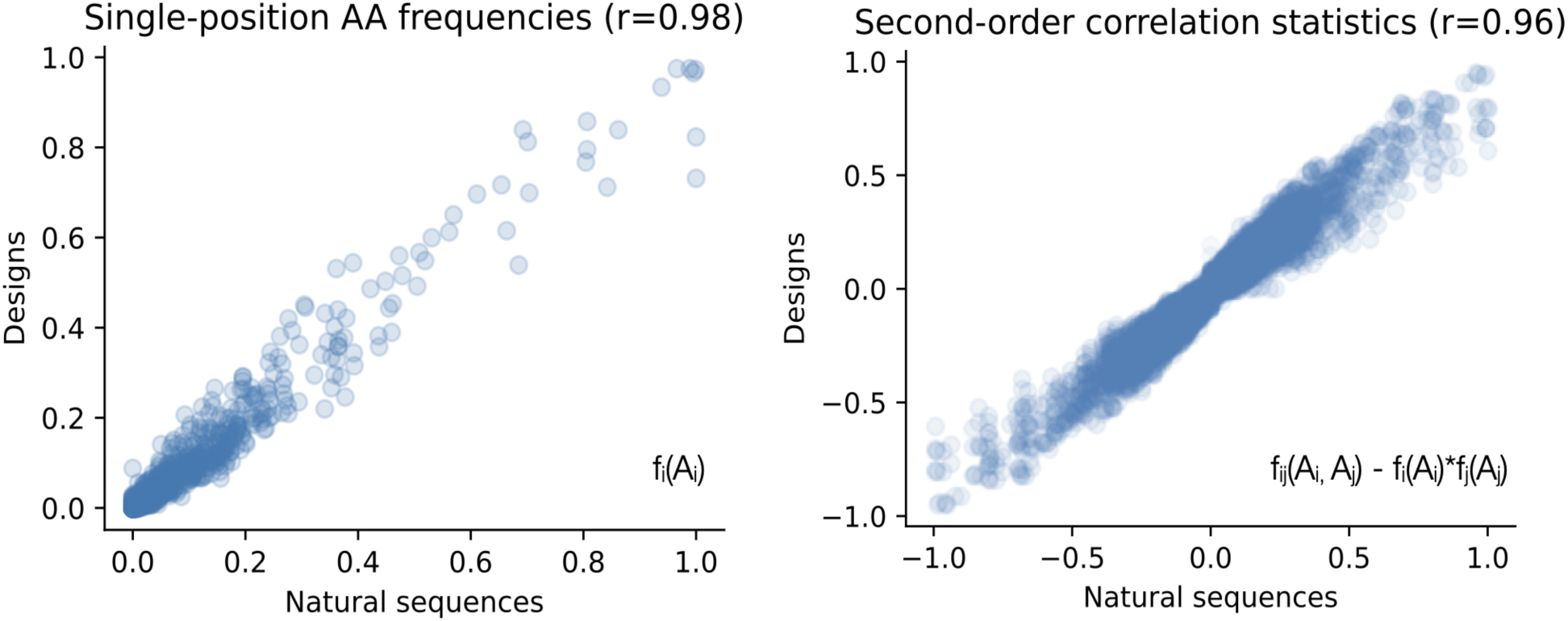
Chorismate mutase designs from EVmutation2 recapitulate statistics of natural sequences. The single-position amino acid frequencies *(left panel)* and pairwise covariance statistics *(right panel)* of sampled chorismate mutase sequences agree with those of natural sequences used to build the EVmutation2 model, indicating that the model captures functional constraints and pair interactions encoded in the sequences.

**Supplementary Figure 6:**
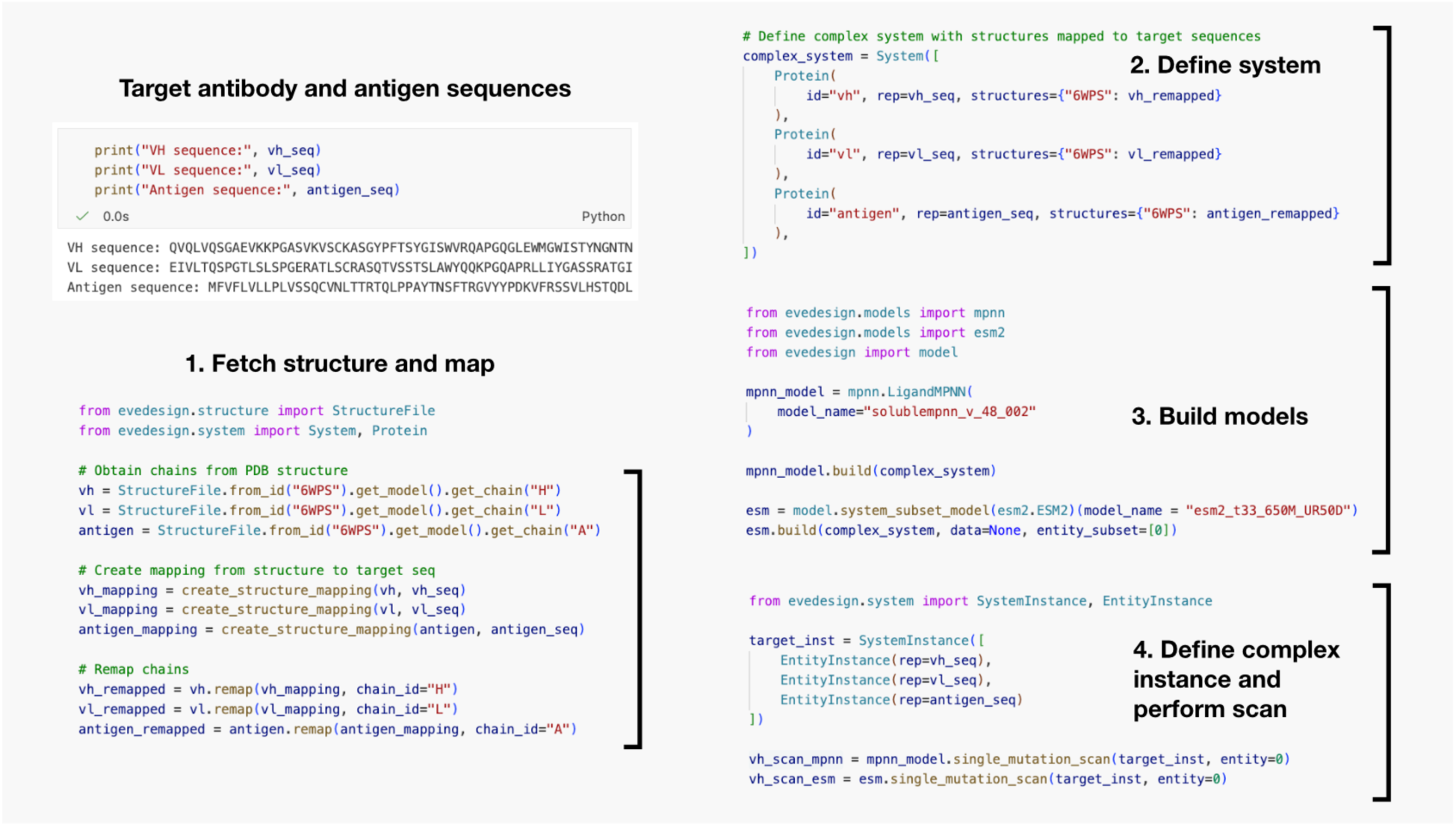
Code demonstration of an antibody maturation pipeline implemented using evedesign. The pipeline implements a sequence-based model (ESM-2) and a structure-based model (ProteinMPNN). The workflow can be summarised into four steps: fetching a complex structure and mapping it to the target sequences, defining the system, building the models, and predicting single substitutions on a complex instance.

**Supplementary Figure 7:**
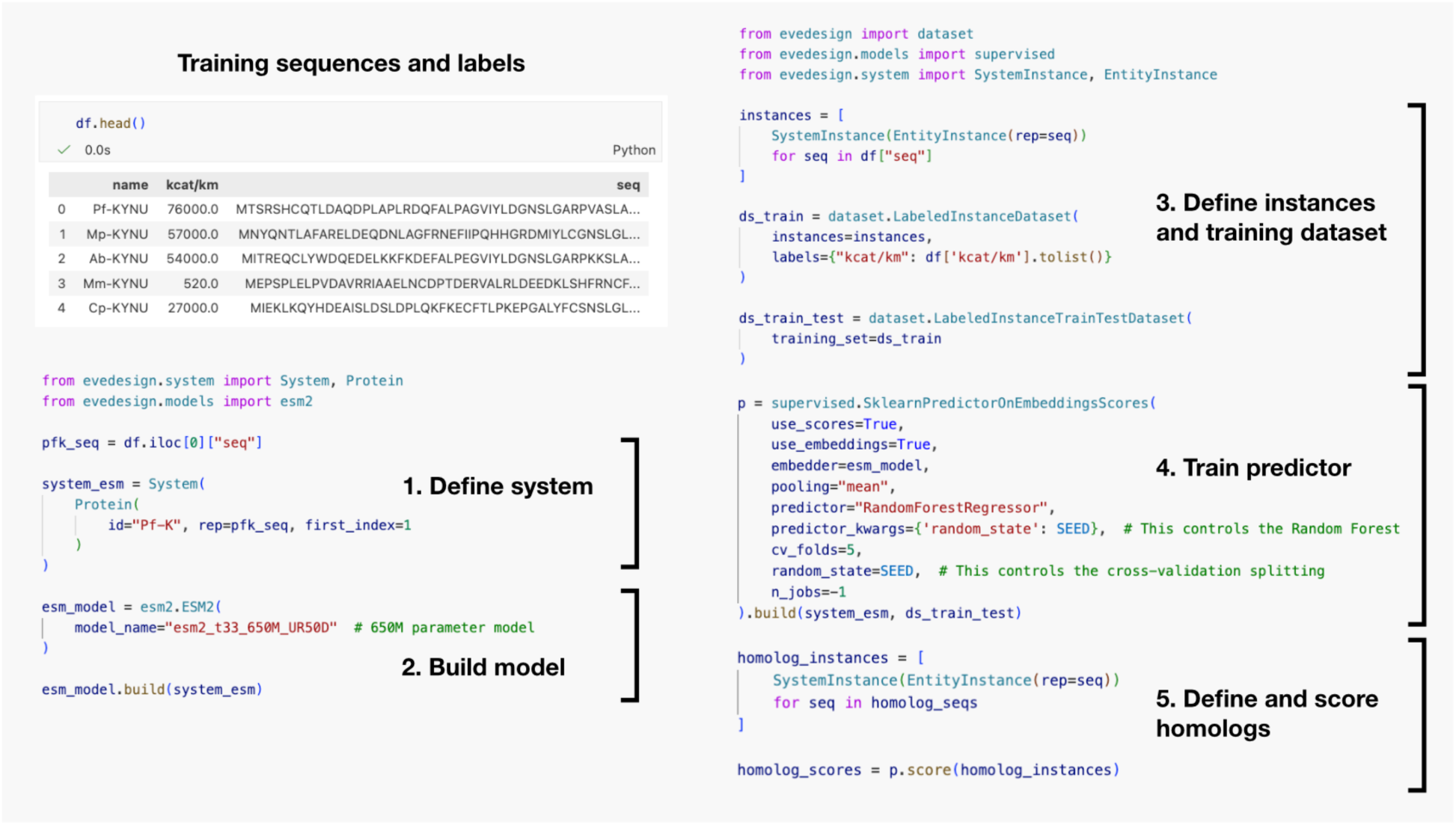
Code demonstration of a supervised evedesign pipeline for predicting catalytic efficiencies of kynureninases. The pipeline can be summarised into five steps: defining the system, building the model, defining training data instances, training the predictor, and defining and scoring homolog instances^57^. A Random Forest is used as the predictor, but any scikit-learn predictor can be used. Model scores are also used as features of the regressor.

**Supplementary Figure 8:**
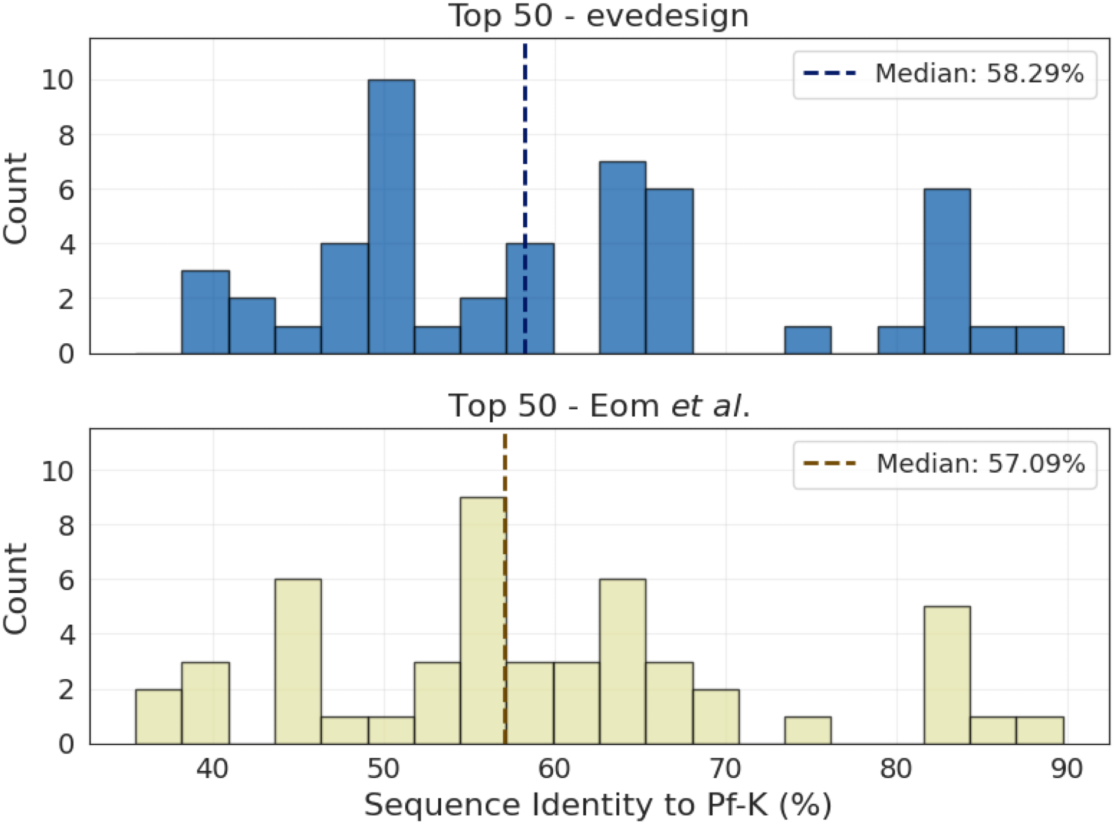
An evedesign-based supervised predictor of catalytic efficiencies of KYNase homologs captures previously reported patterns. The distribution of sequence identities to the starting *Pseudomonas fluorescens* KYNase (Pf-K) of the 50 highest-ranked KYNase homologs obtained with evedesign (*top panel*) and those obtained in Eom *et al.*^57^ (*bottom panel*). Sequence identity ranges and medians are similar. KYNase, kynureninase.

## Notes

### Competing Interest Statement

T.A.H. consults for Seismic Therapeutic and Tectonic Therapeutic, and is a co-founder of Seismic Therapeutic. C.S. is on the SAB of Cytoreason Ltd. M.S. acknowledges outside interest in Stylus Medicine. D.S.M. consults for GenBio.ai, Dyno Therapeutics, Octant, Jura Bio, Tectonic Therapeutic and Genentech, and is a co-founder of Seismic Therapeutic. The remaining authors declare no competing interests.

